# Roosting ecology and the evolution of bat landing maneuvers

**DOI:** 10.1101/2021.10.21.465259

**Authors:** David B. Boerma, Sharon M. Swartz

## Abstract

Biomechanics is poised at the intersection of organismal form, function, and ecology, and forms a practical lens through which to investigate evolutionary linkages among these factors. We conducted the first evolutionary analysis of bat flight dynamics by examining the phylogenetic patterning of landing mechanics. We discovered that bats perform stereotyped maneuvers that are correlated with landing performance quantified as impact force, and that these are linked with roosting ecology, a critical aspect of bat biology. Our findings suggest that bat ancestors performed simple, four-limbed landings, similar to those performed by gliding mammals, and that more complex landings evolved in association with novel roost types. This explicit connection between ecology and biomechanics presents the opportunity to identify traits that are associated with a locomotor behavior of known ecological relevance, thus laying the foundation for a broader understanding of the evolution of flight and wing architecture in this extraordinarily successful mammalian lineage.

## Introduction

Morphologists and biomechanicians often study organismal evolution as a function of three interrelated factors: structure (morphology), function (mechanics or behavior), and context (ecology). Detecting linkages among traits from these categories and discerning where trait shifts correspond with patterns of diversification not only provides evidence of selection but can also point to specific drivers of adaptive radiations, which are a central phenomenon in evolution (Arbour et al., 2019; Burress and Wainwright, 2019; Dakin et al., 2018; Eliason et al., 2020; Muñoz et al., 2018; Stroud and Losos, 2016). Within this framework, many investigations focus on pairwise relationships between two of the three factors: the discipline of functional morphology relates structure to function, whereas the discipline of ecomorphology relates structure with environmental context. These fields reveal both the variety of forms that evolution has produced and the details of how those forms work. Ultimately, however, each field omits the relationship that can be most informative for understanding the *process* of evolution alongside its products: the relationship between biomechanical function and ecological context. If “organismal performance is the primary substrate upon which selection acts, and variation in performance often arises from variation in biomechanics” (Higham et al., 2016), then directly probing the relationship between biomechanical diversity and ecological diversity can point to specific traits that could be targets of selection, and produce testable hypotheses about how form, function, and ecology interact to drive diversification.

Coordinated shifts in form, function, and context are necessary for the evolution of new locomotor modes, such as flight in the lineages that gave rise to bats, birds, insects, and pterosaurs. Although often overlooked, the evolution of flight required not only the evolution of flight *per se*, but also the evolution of landing maneuvers, which transition an animal from moving in air to a standstill; one need only refer to the Greek myth of Daedalus and Icarus to learn that the capacity for flight without the ability to safely land is untenable. For bats and most other flying animals, landing maneuvers also provide access to the structures that constitute their homes, such as roosts, nests, mounds, and hives. They rely on these structures to provide critical functions that extend beyond simply serving as refugia from weather and predators. For example, roost location determines the foraging grounds of many bat species; roosts serve as social spaces that facilitate access to mates, maternal care, and meal sharing; and divergent roost preferences can drive niche partitioning to permit co-occurrence of closely related species (Herrera et al., 2018; Kunz and Lumsden, 2003; Voss et al., 2016; Wilkinson, 1984). Furthermore, roost types vary among bat species and comprise a wide range of natural and human-made structures. These include bare expanses of cave ceiling, crevices and clefts in rock walls, cavities in trees, the voids beneath exfoliating tree bark, within the culms of bamboo, inside the funnels of furled leaves, and even within pitcher plants (see (Kunz and Fenton, 2006) and (Altringham, 2011) for review). Roosting ecology therefore plays an outsized role in defining the environmental mosaic in which bats survive and evolve. Roosting, along with other aspects of bat ecology, such as diet, foraging style, sensory modalities, etc., shapes the behavioral and environmental context that drives changes in form, function, performance, and ultimately diversity (Higham et al., 2016; Schluter, 2009).

Despite the importance of roosting ecology to extant bat diversity, we know little about mechanistic factors that drive roost choices. Measures of biomechanical performance can yield insight into microhabitat preferences (Moore et al., 2017), and for bats, the mechanics of landing maneuvers may be linked to the physical properties of roosts. Specifically, landing dynamics may facilitate access to particular roost types for some species and reduce access for others. Bat landing maneuvers serve two basic functions: 1) body reorientation, and 2) velocity reduction. Body reorientation positions the claws of the foot and/or thumbs to attach to the roost and transitions the bat from a head-forward posture, with the vertebral column approximately parallel to the ground, to the characteristic head-under-heels roosting posture of most species. Velocity reduction modulates the bat’s impact force with the roost and transitions its body from flight, with the center of mass at a non-zero forward velocity, to roosting with the center of mass at rest.

To date, research has identified three landing maneuvers among four species, which are named according to the number of points of contact the bat uses to attach to its landing site upon contact. These maneuvers include two-point landing (both hindlimbs only), and two variants of a four-point landing (both thumbs plus both hindlimbs) (Boerma et al., 2019; Riskin et al., 2009) (Supplemental Videos 1 – 4). Each landing style also involves a characteristic sequence of body rotations, and results in either relatively high or low impact forces normalized to bodyweight. Four-point landings are rotationally simple, primarily involving body pitch, and result in higher impact forces (>3 bodyweights), whereas two-point landings are the most rotationally complex, and result in low peak impact forces (≤1 bodyweight) (Boerma et al., 2019; Riskin et al., 2009). These studies have suggested that landing maneuvers and roosting habits are mechanically linked such that high-impact landings (four-point) are associated with roosting on compliant foliage or vertical surfaces, whereas low-impact landings (two-point) are associated with roosting on stiff horizontal surfaces, such as cave ceilings or tree hollows.

The broad biological importance of roosting ecology and interspecific variation in landing mechanics offers an opportunity to discover how the biomechanical basis of landing performance may underlie how bats take refuge and disperse within their environment. In the present study, we ask three questions relating to landing mechanics, roosting habits, and the potential associations between them: 1) Do previously documented relationships between landing style and impact force remain consistent across a more diverse sample of bats, 2) what is the evolutionary history of bat landing maneuvers, and 3) is landing style linked to roosting ecology? We hypothesized that (i) rotationally complex landing maneuvers would result in lower impact forces than rotationally simple landing maneuvers, across species and body sizes; (ii) rotationally simple, four-point landings are the ancestral condition for bats from which any other style must have evolved; and (iii) landing styles are associated with the physical properties of the roosts to which they provide access. With respect to this final hypothesis, we predicted that four-point landings would be associated with compliant roosts, such as those constructed from foliage, because they could absorb the high impact forces generated by this landing style and because multiple points of contact enhance stability when landing on unstable targets (Boerma et al., 2019; Bonser, 1999; Demes et al., 1995; Riskin et al., 2009). We also predicted that stiff roosts, such as cave ceilings or tree cavities, would be associated with two-point landings because low impact forces could enhance the control and precision of landings and reduce risk of injury when a flying bat decelerates rapidly to attach to a stiff surface.

## Results

### Landing styles across species

We recorded 665 landings from 35 bat species, representing nine families. Of these, 15 species performed two-point landings, 5 performed three-point landings, and 15 performed four-point landings, including *Thyroptera tricolor*, which performed a specialized four-point landing maneuver; see below and Boerma et al., 2019) (Table 1, Supplemental Videos 1 – 4). Overall, landing style was consistent within and among individuals of each species. Notable exceptions include *Artibeus jamaicensis*, which performed two- (29%) and three-point (71%) landings, and *Miniopterus schreibersii*, which performed two- (18%), three- (36%), and four-point (45%) landings. Pteropodid, vespertilionid, and mormoopid species performed four-point landings; emballonurid, rhinolophid and hipposiderid species performed two-point landings; and phyllostomids performed two-, three-, and four-point landings.

**Table 1:** Study taxa, landing style observed, peak landing impact forces, and roosting ecology category. Roost categories are: cavity in standing tree (CST), exposed on standing stree (EST), unmodified foliage (FOL-UF), furled leaf-tubed (FOL-TB), foliage modified into leaf-tents (FOL-LT), termite or ant nests (TAN), rocks and/or caves (R/C), and rock crevices (CREV). Bolded categories indicate those used for comparative analyses; see text for further explanation. *Data from Riskin et al. (2009). ^†^*H. pratti* performed landings that were qualitatively similar to two-point landings, however, following attachment with the hindlimbs, bats flexed the spine ventrally and extended the shoulder and elbow joints to lift the thumb claws ventrally toward their attachment site on the landing plate. Because the thumbs were attached only after the contact during landing, we classify landings by *H. pratti* as two-point in our analysis. ^‡^*T. tricolor* performs a specialized four-point landing maneuver (see Boerma et al., 2019). See Figure 1–Source Data 1 for raw data used to generate this table.

| Taxon | N (total observed landings, individuals) | N (Force recordings, individuals) | Landing Style | Peak $F_{tot}$ (Bodyweight BW) (mean±s.d) | Roosting Ecology | Source(s) |
| --- | --- | --- | --- | --- | --- | --- |
| **<u>Yinterpochiroptera</u>** |  |  |  |  |  |  |
| Pteropodidae |  |  |  |  |  |  |
| *Cynopterus brachyotis* * | 30, 3 | 30, 3 | 4-point | 3.83±1.23 | FOL-UF, **FOL-LT** | (Campbell et al., 2006; Funakoshi and Zubaid, 1997; Tan et al., 1997) |
| *Rousettus aegyptiacus* | 57, 3 | – | 4-point | – | R/C | (Herzig-Straschil and Robinson, 1978; Kwiecinski and Griffiths, 1999; Thomas and Fenton, 1978) |
| Hipposideridae |  |  |  |  |  |  |
| *Hipposideros pratti* | 7, 1 | – | 2-point<sup>†</sup> | – | R/C | (Niu et al., 2007; Zhang et al., 2009)( |
| Rhinolophidae |  |  |  |  |  |  |
| *Rhinolophus hipposideros* | 33, 3 | 32, 3 | 2-point | 0.52±0.04 | R/C | (Aulagnier et al., 2018) |
| *Rhinolophus mehelyi* | 30, 4 | 30, 4 | 2-point | 0.60±0.17 | R/C | (Aulagnier et al., 2018) |
| *Rhinolophus euryale* | 22, 4 | 22, 4 | 2-point | 1.03±0.44 | R/C | (Aulagnier et al., 2018) |
| *Rhinolophus ferrumequinum* | 31, 4 | 31, 4 | 2-point | 1.29±1.00 | R/C | (Aulagnier et al., 2018) |
| **<u>Yangochiroptera</u>** |  |  |  |  |  |  |
| Emballonuridae |  |  |  |  |  |  |
| *Rhynchonycteris naso* | 6, 1 | – | 2-point | – | **EST**, FOL-UF | Sources cited by (Fenton et al., 2001; Voss et al., 2016) |
| *Saccopteryx bilineata* | 2, 1 | – | 2-point | – | **CST**, **EST** | Sources cited by (Voss et al., 2016) |
| Thyropteridae |  |  |  |  |  |  |
| *Thyroptera tricolor* | 71, 16 | 44, 14 | 4-point<sup>‡</sup> | 6.98±1.89 | FOL-TB | (Wilson and Findley, 1977) |
| Mormoopidae |  |  |  |  |  |  |
| *Pteronotus mesoamericanus* | 17, 2 | – | 4-point | – | R/C, CST | Sources cited by (Voss et al., 2016) |
| *Pteronotus davyi* | 11, 2 | – | 4-point | – | R/C | Sources cited by (Fenton et al., 2001) |
| Phyllostomidae |  |  |  |  |  |  |
| *Micronycteris schmidtorum* | 5, 1 | – | 2-point | – | CST | Sources cited by (Voss et al., 2016) |
| *Glossophaga soricina* * | 49, 5 | 49, 5 | 2-point | 0.63±0.11 | CST, R/C | Sources cited by (Voss et al., 2016) |
| *Chrotopterus auritus* | 3, 1 | – | 2-point |  |  |  |
| *Mimon cozumelae* | 20, 4 | 19, 4 | 2-point | 1.22±0.29 | CST, R/C | Sources cited by (Fenton et al., 2001; Simmons and Voss, 1998; Voss et al., 2016) |
| *Lophostoma evotis* | 9, 1 | – | 3-point | – | TAN | (Fenton et al., 2001; Reid, 2009) |
| *Gardnerycteris crenulatum* | 3, 1 | – | 2-point | – | CST | Sources cited by (Voss et al., 2016) |
| *Carollia sowelli* | 16, 3 | 16, 3 | 2-point | 1.98±0.41 | CST, R/C | (Fenton et al., 2001); Sources cited by (Voss et al., 2016) |
| *Carollia perspicillata* * | 50, 5 | 50, 5 | 2-point | 0.76±0.15 | CST, CFT, R/C | (Cloutier and Thomas, 1992; Fenton et al., 2001; Reid, 2009; Voss et al., 2016) |
| *Sturnira parvidens* | 23, 5 | 11, 3 | 4-point | 4.22±2.29 | FOL-UF, CST | (Fenton et al., 2001, 2000) |
| *Uroderma bilobatum* | 2, 1 | 2, 1 | 3-point | 3.95 | FOL-LT | (Barbour, 1932; Timm, 1985) |
| *Dermanura phaeotis* | 25, 5 | 10, 3 | 3-point | 2.73±2.32 | FOL-LT | (Timm, 1985) |
| *Artibeus jamaicensis* | 21, 3 | 21, 3 | 2-, 3-point | 1.19±0.30 | R/C, FOL-LT | Sources cited (Timm, 1985) |
| *Artibeus intermedius* | 11, 2 | 11, 2 | 2-point | 1.03±0.24 | R/C, CST, FOL-UF, FOL-LT | (Reid, 2009) |
| *Artibeus watsoni* | 5, 1 | – | 3-point | – | FOL-LT | (Chapman, 1932; Chaverri and Kunz, 2006; Choe and Timm, 1985) |
| Miniopteridae |  |  |  |  |  |  |
| *Miniopterus schreibersii* | 30, 3 | 11, 2 | 2-, 3-, 4-point | – | R/C, CREV | (Aulagnier et al., 2018; Nowak, 1999) |
| Vespertilionidae |  |  |  |  |  |  |
| *Myotis keaysi* | 5, 1 | – | 4-point | – | R/C, **CREV**,<br>CST | (Brunet and Medellín, 2001;<br>Hernández-Meza et al., 2005;<br>Jr. et al., 1973; Reid, 2009) |
| *Myotis daubentonii* | 10, 1 | – | 4-point | – | **CREV**, R/C | (Aulagnier et al., 2018;<br>Bogdanowicz, 1994) |
| *Myotis myotis* | 12, 2 | – | 4-point | – | R/C, **CREV** | (Aulagnier et al., 2018) |
| *Myotis capaccinii* | 16, 4 | 4, 1 | 4-point | 4.26 | R/C, **CREV** | (Aulagnier et al., 2018;<br>Papadatou et al., 2008) |
| *Rhogeessa aeneus* | 3, 1 | – | 4-point | – | CST, **CREV**,<br>FOL-UF | (Nowak, 1999; Reid, 2009) |
| *Eptesicus fuscus* | 10, 1 | 9, 1 | 4-point | 5.46 | **CREV**, CST | (Brigham, 1991; Lausen and<br>Barclay, 2003, 2002) |
| *Eptesicus serotinus* | 10, 1 | – | 4-point | – | **CREV**, R/C | (Aulagnier et al., 2018; Catto<br>et al., 1995) |
| *Hypsugo savii* | 10, 1 | – | 4-point | – | **CREV**, R/C | (Aulagnier et al., 2018;<br>Horáček and Benda, 2004) |
| **Totals:** | **665, 96** | **401, 65** |  |  |  |  |

### Landing impact force increases with points of contact across landing styles

Two-point landings from 32 individuals of 15 species, from 4 families, uniformly resulted in low peak impact forces, with a mean of 0.95 ± 0.54 BW (mean ± s.d.,). Four-point landings resulted in higher impact forces: 3.72 ± 1.71 BW (n=10 individuals, 5 species). Three-point landings were intermediate in magnitude and more variable; mean impact was 1.71 ± 1.44 BW (n=12 individuals, 4 species). The specialized four-point landings of *T. tricolor* resulted in the highest impact forces, 6.98±1.89 BW (n=14 individuals) (Boerma et al., 2019). Phylogenetic generalized least squares regression (PGLS, *T. tricolor* omitted, see *Phylogenetic Analyses* in Methods) revealed that log peak impact force increases significantly with points of contact across species (DF=14, F=33.47, p=4.726 × 10^−5^). Phylogenetic ANOVA (*T. tricolor* omitted) corroborated that landing style has a significant effect on log peak impact force (F=14.04, p=0.0099). Pairwise posthoc tests with Holm-Bonferroni correction show that two-point landings result in significantly lower impact forces than four-point landings (t= -5.26, p=0.0078), and that the intermediate impact forces associated with three-point landings are not statistically different from either two-point (t= 2.49, p=0.2266) or four-point landings (t=-2.45, p=0.2266) (see Source Code File 3 and PGLS phylANOVA–Source Data 1 for the raw data and code used to conduct these analyses).

### Four-point landings are ancestral and preceded multiple independent evolutions of two- and three- point landings

We simulated 1000 stochastic character maps of landing style on a phylogeny pruned to our sampled taxa (figure 2). These simulations estimated that four-point landings were ancestral (Posterior Probability (PP) = 0.862), and that landing style shifted an average of 7.903 times. Of these shifts, 3.161 state changes occurred from four- to two-point landings. This was the most common evolutionary shift and occurred at multiple locations in the bat phylogeny. Additional state changes were concentrated among bats in the family Phyllostomidae. In this clade, 2.061 shifts occurred from two- to three-point landings, and we detected 1.427 reversals from two- to four-point landings (in *S. parvidens*). Our reconstruction also estimated 0.636 state changes from three- to four-point landings, 0.384 state changes from four- to three-point landings, and 0.234 state changes from three- to two-point landings. The mean proportion of time spent in each state was 54.14% in four-point landings, 38.90% in two-point landings, and 7.06% in three-point landings.

Among the taxa we sampled, we detected three independent shifts to from four- to two-point landings. These occurred at the base of the clade giving rise to the Rhinolophidae and Hipposideridae (PP_two-point_ = 0.642), at the base of Emballonuridae (PP_two-point_ = 0.92), and (iii) at the base of the Phyllostomidae (PP_two-point_ = 0.949) (figure 2A, B, & C). Three-point landings evolved relatively recently, emerging first in the common ancestor of the phyllostomid subfamily Stenodermatinae (figure 2F). This ancestor possessed equal probability of performing two- or three-point landings (PP_two-point_ = 0.464; PP_three-point_ = 0.463) (figure 2F). In this species sample, the common ancestor of tent-roosting phyllostomids (stenodermatines excluding *S. parvidens*) was most likely to perform three-point landings (PP_three-point_ = 0.921). Three-point landings also arose in the phyllostomid *L. evotis* (figure 2D). We detected one reversal from two- to four-point landings in the phyllostomid *S. parvidens* (figure 2E). Four-point landings, the ancestral condition, persisted in pteropodids, mormoopids, and vespertilionids, based on analysis of this sample.

### Landing styles are associated with the physical properties of roosts across species

We investigated the relationship between roosting ecology and landing style by using phylogenetic logistic regression to compare landing style with roosting ecology using alternative roost classification schemes, which aggregated roost categories with similar physical characteristics (table 2). Compared to the null model, our aggregated model had greater explanatory power for predicting landing style from roosting ecology, as indicated by AIC score. Our null model, which tested for association between roosting habits and landing style using the most common roost type for each species, revealed a significant positive association only between cavity-roosting and two-point landings (ß_NullCST-2pt_ = 3.862; p_Null,CST-2pt_ = 0.01701). Aggregating roosting categories according to physical properties, such as compliance, orientation, and spatial constraint, allowed us to test the hypothesis that these physical properties are significantly associated with the mechanics of the three known landing styles. We found that two-point landings were positively associated with stiff, horizontal roosts, such as caves and cavities (ß_Agg,[CST+EST+R/C]-2pt_ = 4.367; p_Agg,[CST+EST+R/C]-2pt_ = 0.008089). Three-point landings were positively associated with roosting in spatially constrained structures, such as leaf-tents and termite nests (ß_Agg,[tent+tan]-3pt_ = 3.525; p_Agg,[tent+tan]-3pt_ = 0.04354). Four-point landings were negatively associated with roosting beneath stiff, horizontal structures (ß_Agg,[CST+EST+R/C]-4pt_ = -2.144; p_Agg,[CST+EST+R/C]-4pt_ = 0.03589).

**Table 2:** Correlations between landing style and roosting ecology from phylogenetic logistic regressions. We provide Firth-corrected coefficient estimates (ß) with bootstrapped 95% confidence intervals (in brackets) and Wald p-values (in italics) to denote significant associations between roost type. Significant p-values are bolded and set within shaded cells. AIC scores provide comparison between our Null and Aggregated model (models with smaller AIC are preferred; differences are meaningful when ≥ 2). P-values are conditional upon phylogenetic signal, α, where values near 0 denote strong phylogenetic signal and values approaching 1 indicate weak phylogenetic signal. Roosting ecology categories correspond with those listed in Table 1: cavity in standing tree (CST), exposed on standing tree (EST), rocks and/or caves (R/C), termite or ant nests (TAN), foliage-leaf tent (FOL-LT), unmodified foliage (FOL-UF), and rock crevices (CREV). See Table 2–Source Data 1 and Source Code File 3 for data and code used to produce this table.

| Model | Landing | CST | EST | R/C | TAN | FOL-LT | FOL-UF | CREV | AIC | $\alpha$ |
| --- | --- | --- | --- | --- | --- | --- | --- | --- | --- | --- |
| | | $\beta$= Mean coefficient estimate, [lower CI, upper CI], *pval* | | | | | | | | |
| All Roost Types (Null) | 2-pt. | 3.862<br>[0.637, 5.413]<br>*0.01701* | 2.944<br>[-0.053, 3.700]<br>0.26658 | 1.978<br>[-1.225, 4.072]<br>0.12250 | 1.748<br>[-0.869, 3.164]<br>0.2981 | 0.594<br>[-2.345, 2.482]<br>0.68152 | 1.027<br>[-1.225, 2.655]<br>0.50338 | -1.277<br>[-2.737, 0.216]<br>0.30867 | 40.61 | 0.0036 |
|  | 3-pt. | -0.098<br>[-1.708, 2.136]<br>0.8973 | -0.135<br>[1.099, 1.408]<br>0.9097 | -0.975<br>[-2.402, 0.471]<br>0.2719 | 2.459<br>[-0.580, 3.340]<br>0.4405 | 1.608<br>[-0.524, 4.513]<br>0.3182 | 0.090<br>[-2.402, 0.471]<br>0.2719 | -0.453<br>[-1.243, 0.680]<br>0.7954 | 30.09 | 0.0030 |
|  | 4-pt. | -0.657<br>[-2.108, 0.557]<br>0.9017 | -0.237<br>[-1.627, -1.315]<br>0.8468 | -0.237<br>[-1.812, 1.878]<br>0.7521 | -0.234<br>[-1.535, 1.122]<br>0.8523 | -0.237<br>[-1.275, 1.063]<br>0.7938 | 1.081<br>[-0.995, 3.199]<br>0.5321 | 0.190<br>[-1.020, 1.090]<br>0.9017 | 42.35 | 0.0045 |
| Aggregated Model (Agg) | 2-pt. | 4.367<br>[2.023, 5.428]<br>*0.008089* |  |  | 1.648<br>[-0.851, 3.065]<br>0.366708 |  | 1.346<br>[-0.519, 3.345]<br>0.580319 | -2.947<br>[-3.023, -1.079]<br>0.366708 | 33.03 | 0.5251 |
|  | 3-pt. | -0.631<br>[-2.476, 1.681]<br>0.76677 |  |  | 3.525<br>[0.231, 5.184]<br>*0.04354* |  | 1.356<br>[-0.532, 3.348]<br>0.57546 | -2.937<br>[-3.00, -0.248]<br>0.05402 | 20.28 | 0.9100 |
|  | 4-pt. | -2.144<br>[-4.498, -0.337]<br>*0.03589* |  |  | -1.133<br>[-3.681, 1.890]<br>0.38111 |  | 1.308<br>[-2.236, 2.538]<br>0.56684 | 0.703<br>[-0.637, 2.950]<br>0.51964 | 32.33 | 0.0076 |

## Discussion

Using a combination of field and lab-based measurements, we investigated functional links between landing mechanics and roosting ecology, which is a critical biological factor for bats. Our measurements of landing style in 35 bat species and peak impact forces in a 17 species subset of this group shows that landing impact force increases with the number of points of contact a bat uses to land, i.e. impact force varies according to landing style even after correction for phylogenetic relationships among the study species. Moreover, we observe that bat landing styles are associated with patterns of roost use: rotationally simple, high-impact four-point landings are ancestral for bats, and rotationally complex two-point landings evolved independently multiple times in lineages that habitually roost beneath stiff surfaces. Furthermore, in the stenodermatines, a subfamily of the Phyllostomidae that shows a reversal from roosting in cavities (stiff surfaces) to roosting in foliage (compliant surfaces) (Garbino and Tavares, 2018), we observe a concomitant reversal from low-impact two-point landings to higher-impact three- and four-point landings. Three-point landings, which we describe for the first time in the present study, arose twice among our sampled taxa, each time in species that roost within spatially constrained horizontal roosts, such as leaf tents or evacuated termite nests.

### Roosting ecology and the evolution of bat landing maneuvers

Four-point landings, the ancestral condition for bat landings, are performed by nearly half of the species in our sample (15 of 35). These landings were negatively associated with roosting beneath stiff, horizontal surfaces (e.g., tree cavities and cave ceilings), but are not strictly associated with compliant foliage roosts across the bat phylogeny, as has been hypothesized in earlier investigations (Riskin et al., 2009). This foliage-roost hypothesis is weakly supported in the phyllostomids we examined (e.g., *S. parvidens*) and for *T. tricolor*, both of which are foliage roosting species that employ four-point landings. However, our sampling of pteropodids, mormoopids, and vespertilionids, which included bats that habitually roost beneath stiff surfaces, such as cave ceilings, and those that land on vertical walls and roost within rock crevices (vespertilionids), did not show a clear correlation between four-point landing and foliage roosting. Broader sampling among pteropodids and mormoopids could reveal additional patterns of roost use and landing mechanics. However, our results suggest that landing maneuvers in these three families are not as labile as in phyllostomids or rhinolophids. In the case of crevice-roosting bats, all of which are vespertilionids in this sample, four-point landings may offer functional opportunities despite the higher impact forces typically incurred on stiff substrates. These include facilitating more rapid access to interstices in the walls of cliffs, caves, trees, and human-made structures compared to other landing styles. Landing with four points of contact immediately places all limbs on the substrate, thus allowing for immediate transition from flight to landing to terrestrial locomotion (crawling) along the roost surface. This rapid locomotor transition could minimize the time required to locate crevice refuges (Supplementary Video 5, *M. myotis*) and reduce exposure to predators or adverse climatic conditions.

We observed convergent shifts from four-point to two-point landings at three nodes in the phylogeny (figure 2A, B, and C), each representing a common ancestor of a lineage characterized by roosting beneath stiff, horizontal roosts (table 2). These shifts support the hypothesis that rotationally complex, low-impact two-point landings evolved in association with the physical properties of roosts in these lineages. Further support for this hypothesis is found among the phyllostomids in particular, in which secondary reversals away from stiff horizontal roosts (e.g., cavity or cave-roosting) to roosting in foliage or within spatially constrained structures on vegetation (e.g., leaf-tents and abandoned termite nests), corresponded with shifts from two-point to four- or three-point landings (figure 2D, E, and F). Apart from *L. evotis*, which roosts in termite nests, we documented high-impact three- and four-point landings only among bats in the phyllostomid subfamily Stenodermatinae (figure 2, taxa highlighted by the green vertical line), the lineage in which foliage roosting re-arose within Phyllostomidae (Garbino and Tavares, 2018). If bat landing maneuvers are adapted to the physical properties of roosts, this transition from low-impact to high-impact landings at the node with a corresponding shift from stiff to compliant roosts could signal relaxed selective pressure for low-impact landings.

The stenodermatines are a relatively recent radiation (Amador et al., 2018; Rojas et al., 2016; Shi and Rabosky, 2015), and they show diversification rates that are approximately twice as high those background rates for Chiroptera (Dumont et al., 2012; Shi and Rabosky, 2015). Previous work has identified shifts in diet, sensory modalities, and associated cranial morphology as key innovations that led to this rapid diversification (Arbour et al., 2019; Dumont et al., 2014, 2012; Santana et al., 2012), but some have speculated that shifts toward foliage roosting may have also contributed to increased speciation rates in this clade (Garbino and Tavares, 2018; Voss et al., 2016). Here, we document transitions in this lineage from two-point to three- and four-point landings, and thus hypothesize that these evolutionary shifts in landing mechanics could be included among the factors contributing to the recent evolutionary success of the stenodermatines.

### Many-to-one mapping of high-impact landings

The hypothesis that convergence in roosting habits is associated with convergence of landing style across the bat phylogeny is implicit in our prediction that roosting ecology and landing style are linked. Our findings largely supported this hypothesis, but one intriguing example of where results diverged from this pattern is in lineages that convergently evolved a highly derived roosting ecology – tent-making. Tent-making refers to a behavior in which bats weaken the veins of large leaves by biting them so that portions of the leaf droop to create a tent-like shelter (Barbour, 1932; Kunz and Fenton, 2006; Kunz and Lumsden, 2003; Tan et al., 1997; Timm, 1987). Leaf tents can take multiple forms (see Kunz and Lumsden 2003 for review), but seem to function primarily as refugia from climate, rather than from predators (Tan et al., 1997). This behavior independently arose in at least three species in the family Pteropodidae (represented by *C. brachyotis* in our sample), and several species in the family Phyllostomidae (subfamily Stenodermatinae, figure 2) (Kunz and Lumsden, 2003). Among the species in our study that convergently evolved this derived roosting ecology, we observed family-level differences in landing maneuvers, including the number of points of contact (three vs. four) and limb contact order (hindlimbs first in three-point landings vs. thumbs first in four-point landings). Despite these differences, however, three-point and four-point landings share similar degrees of rotational complexity and result in similarly high impact forces (Figure 1).

**Figure 1:**
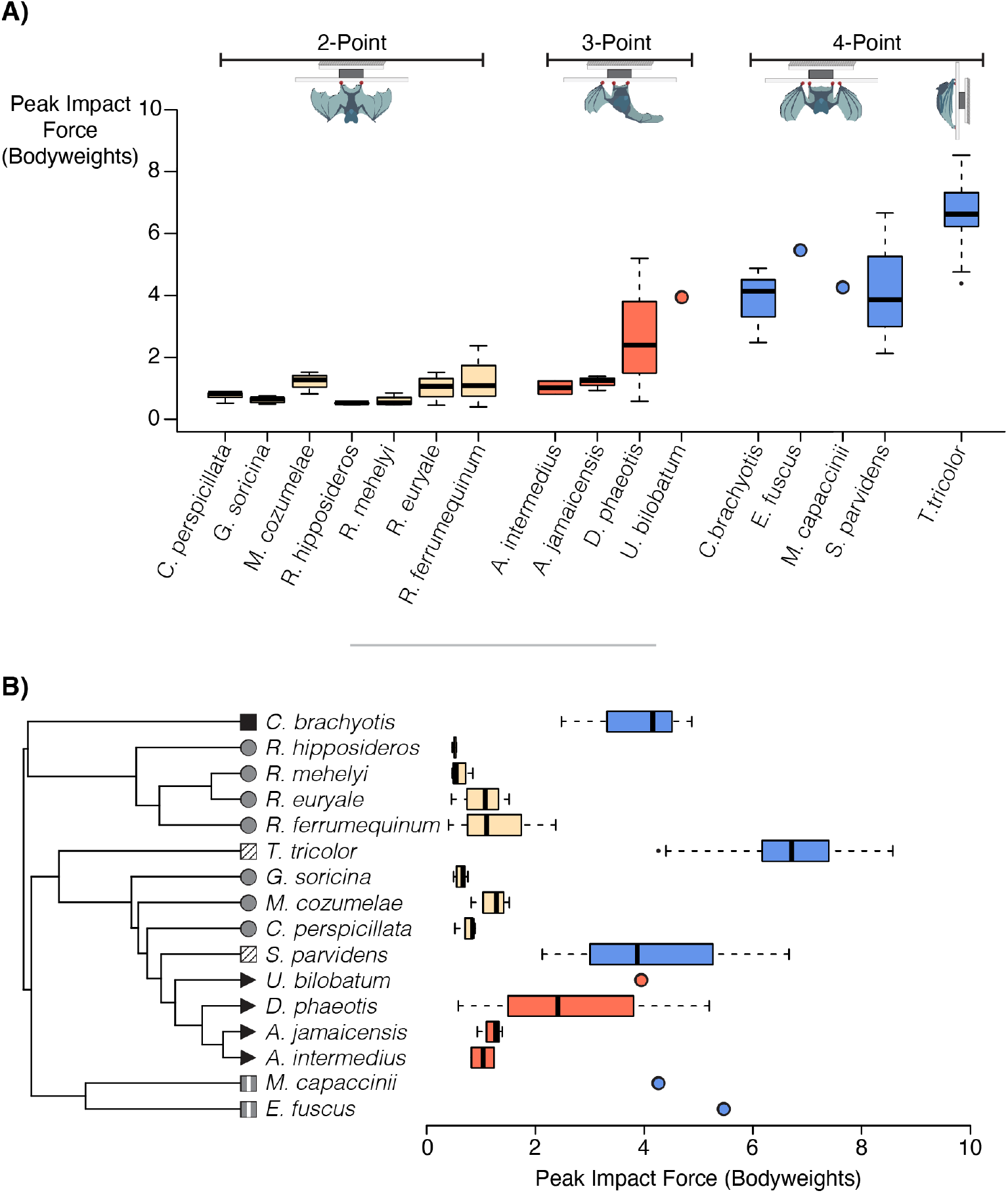
Peak landing impact forces. (excluding *M. schreibersii*; see table 1 for sample sizes). Box and whisker plots show the median and interquartile range. (A) Landing impact forces arranged by landing style. (B) Landing impact forces arranged phylogenetically (tree adapted from Shi and Rabosky, 2015). Two-point landings are denoted by yellow boxes and circle icons at branch tips, three-point landings by red boxes and triangle icons, and four-point landings by blue boxes and square icons. Icon fill color represents roosting ecology: solid gray = stiff horizontal roosts; grey with vertical white stripe = crevices; black = leaf tents; and hatched = unmodified foliage. Legend also provided in figure 2. See Figure 1–Source Data 1 for raw impact force measurements for each landing and Figure 1–Source Data 2 for each individual’s mean peak impact force, the latter of which was used to generate these figures. See Source Code File 1 for R code to reproduce these plots.

**Figure 2:**
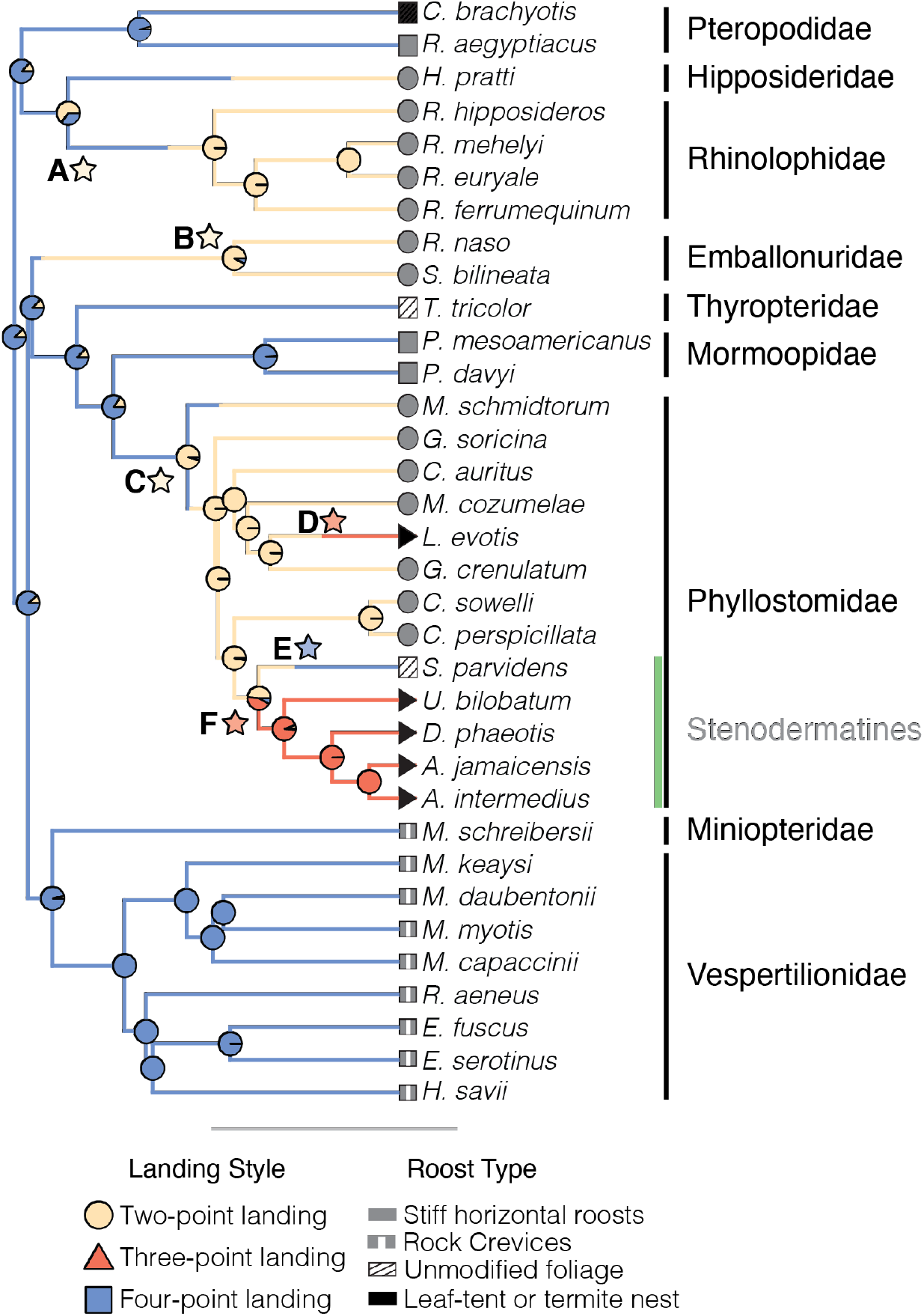
Stochastic map of landing styles. Pie charts at the nodes show posterior probabilities. Stars A-E mark shifts in landing style. Tip shapes denote landing style, and tip fill denotes the roost type used in the aggregated model of phylogenetic logistic regression. Black vertical bars to the right of the species names denote families; the green line highlights the subfamily Stenodermatinae. Phylogeny adapted from Shi and Rabosky (2015). See Figure 2–Source Data 1, Figure 2–Source Data 2, and Source Code File 2 for the raw data and code used to generate this figure. Full posterior probabilities are provided in Supplemental Files 1 and 2.

**Figure 3.**
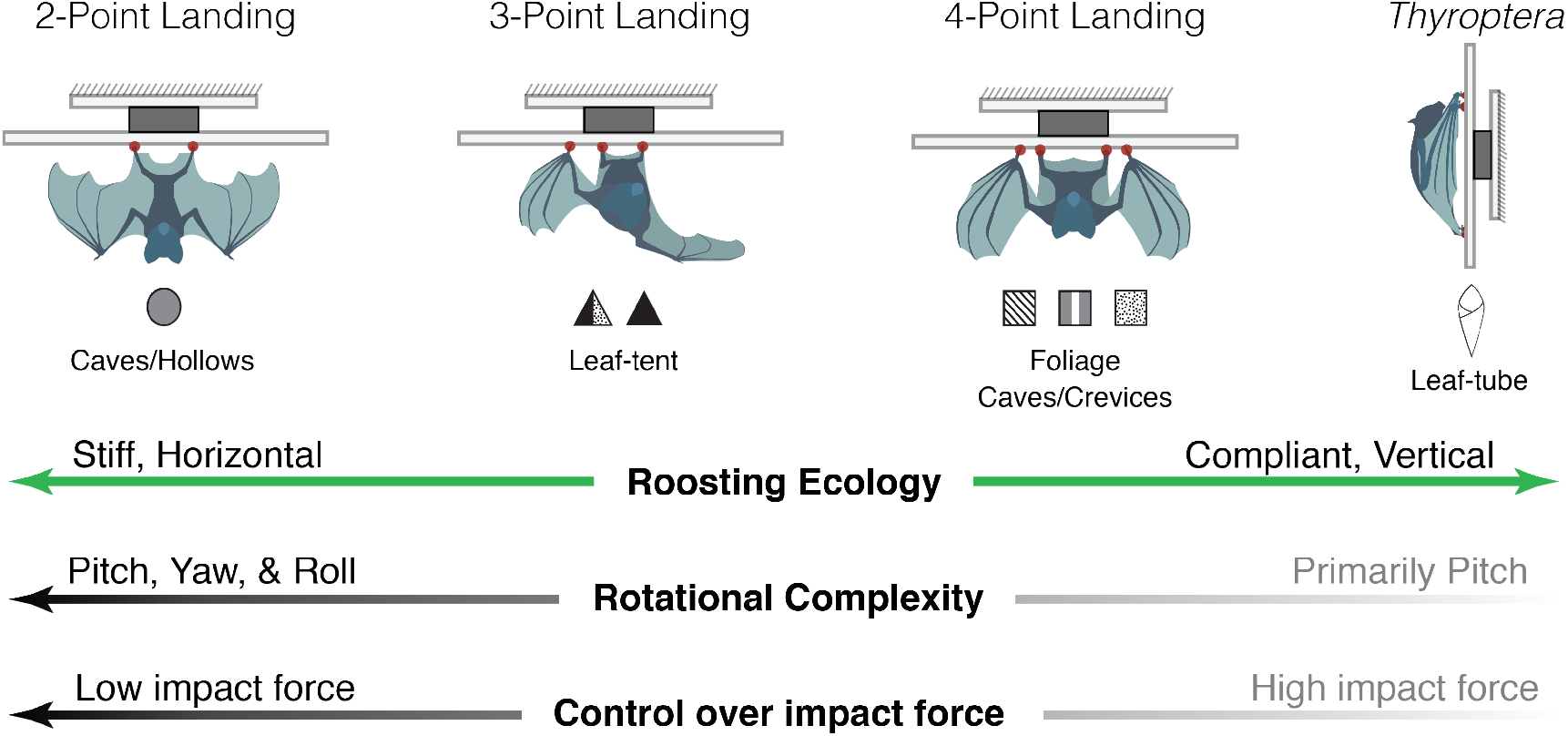
Continuum of landing style, roosting ecology, and landing mechanics. The mechanics of bat landings correspond with patterns of roost use among sampled bats.

This observation suggests a many-to-one mapping of landing mechanics to landing impact force for species that roost in leaf tents; that is, although three- and four-point landings differ kinematically, they result in a similar functional outcome (high impact forces), and their differences may be due simply to the different evolutionary starting points of extant pteropodids and stenodermatines (Wainwright et al., 2005). The most recent common ancestor of stenodermatines and other phyllostomids in our sample most likely roosted in cavities (Garbino and Tavares, 2018) and performed two-point landings (PP_two-point_ = 0.978). Thus, this common ancestor likely landed with low impact force, using only the two hindlimbs as points of contact. Under our hypothesis, the transition to roosting in compliant leaf-tents would have reduced the selective pressure on low-impact landings, thus permitting a shift to higher-impact three-point landings that retained the feet-first contact order but added the thumb as a stabilizing point of contact. In contrast, the common ancestor of the pteropodids (which include *C. brachyotis*) performed four-point landings (PP_four-point_ = 0.95), a landing style already amenable to roosting in compliant leaf tents which can absorb the high-impact landings.

### Other factors that may influence landing style

We focused on associations between landing mechanics and roosting ecology in the present study, but other traits could also influence diversity of landing maneuvers among bats. Here we highlight a couple, including sensory ecology and wing morphology.

A bat’s ability to sense the location, geometry, and surface characteristics of a potential landing site contributes to its capacity to execute accurate, precise landings. Therefore, variation in sensory ecology, specifically echolocation capacity and call structure, could also influence the landing maneuvers of bats. Most bats navigate their environments and detect prey using laryngeal echolocation, and the structure of these echolocation calls, including amplitude, frequency, and rate, differs among species and tasks in ways that trade off between target resolution and detection distance (Geva-Sagiv et al., 2015; Schnitzler et al., 2003). Pteropodids are a notable exception, however, and rely either on vision or rudimentary forms of echolocation such as tongue or wing clicks (Boonman et al., 2014; Jones and Teeling, 2006; Kulzer, 1956; Teeling, 2009). Most studies of echolocation have focused on its role in prey capture or navigation through the environment during forward flight. Little work to date has investigated echolocation behavior when approaching stationary targets, such as roosts (but see (Tian and Schnitzler, 1997)) Interspecific variation in echolocation behavior during landing could reveal patterns that coincide with differences in impact forces and body rotations as bats call to sense the roost during approach. Additionally, in the Pteropodidae, which do not possess laryngeal echolocation, landing behavior could be constrained due to sensory limitations in their capacity to resolve details of potential roosts with high temporal resolution during an approach flight.

Interspecific differences in wing morphology may also relate to variation in landing mechanics. Because aerodynamic forces are highly dependent upon the velocity of airflow over the wings, and landing occurs at low speeds, bats accomplish landing maneuvers using inertial forces almost exclusively (Bergou et al., 2015). The wing’s capacity to effect body rotation via inertial torques is therefore related to their mass moment of inertia, which in turn is determined by the distribution of mass within the wings. Studies that characterize interspecific differences in wing mass distribution may therefore reveal a relationship between the wing’s body mass-normalized mass moment of inertia and the rotational complexity of landing maneuvers. For example, variation in wing inertia could arise from interspecific differences in wing length or relative mass of the bones, muscle and skin that comprise the wing, particularly in the distal regions.

### Estimated ancestral landing mechanics provide support for a gliding bat ancestor

In bats, a group for which origins of powered flight in remain unresolved, studying the evolutionary history of landing mechanics provides a complementary perspective to studying the evolution of flight itself. Despite a lack of fossil bat ancestors, most paleontological and biomechanical investigations point to a gliding origin of bat flight. The hypothesized early bat ancestor was likely arboreal, possessed gliding membranes made of skin, and is hypothesized to have performed gliding locomotion similar to that observed in extant mammalian gliders (Bishop, 2008; Curet et al., 2012; Gunnell and Simmons, 2012; Simmons et al., 2008).

If bat flight has its origins in gliding locomotion, then we would expect that ancestral bats might have landed similarly to extant gliding mammals. Mammalian gliders execute landings that rely almost exclusively on pitching rotations and result in high-impact forces (Bahlman et al., 2012; Bishop, 2007, 2006; Byrnes et al., 2008; Paskins et al., 2007). The four-point landings observed in extant bats are a plausible next step for landing maneuvers because they would require only addition of further pitching to the basic glider landing pattern to facilitate landing on the underside of roosts instead of on the vertical side of tree trunks. Indeed, stochastic character mapping provides evidence that the common ancestor of bats performed a four-point landing maneuver, which relies chiefly on pitching rotations with negligible contributions from yaw and roll, and is characterized by high peak impact forces (Riskin et al., 2009). Our biomechanical study of landing therefore provides additional support for the gliding origin of flight in bats.

### Broader implications: bat conservation and adaptive radiation

Like most of the earth’s biodiversity, bats are vulnerable to human disturbance, whether it be through anthropogenic climate change or more proximate issues, such as deforestation, both of which affect the availability and quality of roosts. If landing mechanics are associated with roosting habits then they may affect the extent to which certain species are robust to displacement via roost destruction. Bats with highly specialized roosting ecologies are generally at higher risk for extinction and are less prevalent in disturbed forest fragments (Herrera et al., 2018; Sagot and Chaverri, 2015). In addition to the difficulties associated with locating suitable alternatives, species with specialized landing maneuvers, such as *Thyroptera tricolor* (Boerma et al., 2019), may also encounter a biomechanical barrier to establishing new roosts. In certain cases, the mechanics of bat landing maneuvers may thus mediate roost access by prohibiting certain species from successfully landing on new surfaces if displaced. Conversely, species whose landing styles are more flexible and permissive, or those which are able to perform multiple landing styles, such as *A. jamaicensis* and *M. schreibersii*, may be able to roost more easily on a diverse array of surfaces and thus might be more robust to habitat destruction and deforestation due to anthropogenic intervention and climate change. Analyses that probe the relationship between number of roost types used, landing style, and habitat range are among the future efforts that could help evaluate this hypothesis.

Additionally, studies that integrate biomechanics with ecology and evolutionary history have the potential to reveal key morphological or behavioral innovations that changed the way lineages interacted with their environments and helped to drive adaptive radiations (Burress et al., 2020; Burress and Wainwright, 2019; Muñoz, 2019; Muñoz et al., 2018; Muñoz and Price, 2019; Stroud and Losos, 2016). Here, we suggest that roosting ecology and landing mechanics are functionally linked, and given the broad biological importance of roosting for bat diversity, the potential for landing mechanics to be a mediating factor during the evolution of diverse roosting habits makes this a promising system for studying how ecological opportunity (roosting ecology), form (wing morphology), and function (landing mechanics) interacted over the course of diversification in bats. The extent to which these factors acted as drivers of speciation in certain lineages is unclear, but the present study serves as a foundation for future inquiry these evolutionary relationships.

Such future work would benefit first and foremost from increased sampling, both in terms of phylogenetic breadth and in the number of individuals per species. Sampling bats from the twelve families absent from our sample and pursuing additional sampling in those we included, especially bats with larger body size (>200 g) or specialized roosting ecologies, would strengthen inferences about the evolutionary history of landing mechanics and better resolve correlations between landing style and roosting ecology. Sampling more individuals per species would also provide a better understanding of the levels of intraspecific variation in impact force, landing style, and landing kinematics. However, we acknowledge that the difficulties of field-based biomechanics research may pose challenges for observing landing behavior in a wider array of species. Approaches that eliminate the measurement of impact forces will allow for broader sampling with videography because some species are difficult or impossible to train to land on a small force platform. Furthermore, recording landing videos at known roost locations rather than with captured individuals in a field-based flight arena might also permit broader sample while simultaneously documenting variations in landing style on natural roosts.

In addition to increased sampling, future work would also benefit from efforts to measure and reconstruct the evolutionary history of morphological traits related to landing maneuvers, such as wing mass distribution, which determines the inertial torques bats use to execute landings (Bergou et al., 2015), and other skeletal features relating to limb stresses and landing impact forces. Taken together, these efforts would determine whether there are clade-based links among roosting habits, landing style, wing morphology, and diversification rates. If shifts in roosting ecology were associated with speciation in certain lineages (e.g., stenodermatines), and if roosting ecology, landing style, and wing morphology were linked, then one should detect significant shifts in diversification rates for clades that arise following coordinated shifts in roosting habits and landing style.

## Conclusions

Resolving the connections among form (morphology), function (mechanics), and environmental context (ecology) are central to understanding the evolutionary history of organisms. While form-function (functional morphology) and form-environment (ecomorphology) relationships are often the focus of evolutionary studies, determining linkages between mechanics and ecology are equally critical to understanding how morphological and ecological variation interact with the organismal performance on which selection acts. We have presented the first evolutionary analysis of any aspect of flight dynamics in bats that links specific traits associated with flight performance to a particular aspect of bat ecology. Our survey of landing mechanics across a broad sample of bats revealed that interspecific variation in landing styles varies along a mechanical continuum of rotational complexity and landing impact force, and that the physical properties of bat roosts are associated with particular landing styles. Independent of phylogenetic relationships, rotationally complex, low-impact landings (two-point) were positively associated with stiff, horizontal roosts, whereas rotationally simple, higher-impact landings (three-or four-point) were negatively associated with stiff roosts, and in some cases were positively associated with roosting in compliant foliage or spatially constrained roosts in vegetation. These results highlight the evolutionary interactions between locomotor mechanics and ecology, establish functional links between landing mechanics and roosting ecology in bats, and suggest that these interactions may be a factor both for mediating roost use and for driving diversification in certain clades. By connecting roosting ecology to the biomechanics of landing, we now have the potential to identify traits that are specifically associated with a particular form of locomotor behavior of known ecological relevance. This accomplishment lays the foundation for a broader understand of the evolution of flight and wing architecture in this extraordinarily successful lineage of mammals. To this end, future work should examine additional ecological and morphological correlates and incorporate evolutionary rate analyses to better resolve how landing mechanics and roosting ecology, and other traits interacted throughout bat evolution.

## Materials and Methods

### Focal taxa, field sites, and animal capture

We recorded 665 landings from 96 bats, representing 35 species, and 9 families (table1). We collected all measurements from wild-caught bats except for *Rousettus aegyptiacus* and taxa from Riskin et al. (2009) (*C. perspicillata, G. soricina*, and C. *brachyotis*), which were captive-bred. Our field sites were located in Lamanai, Orange Walk, Belize (Lamanai Outpost Lodge); Barú, Puntarenas, Costa Rica (Haciénda Barú Biological Research Station); Tabachka, Bulgaria (Siemers Bat Research Station, Max Planck Institute); and Shandong, China (Shandong University). We captured bats using mist-netting, hand-netting, and harp traps.

### Landing experiments

At each field site, we observed bat landings within a temporary flight corridor (3 × 1.5 × 2 m) (length × width × height). For all bats except *T. tricolor* (see Boerma et al. 2019), we covered the walls and ceiling with smooth plastic sheeting to prevent bats from landing anywhere but on a ceiling-mounted landing platform, which was covered with stiff plastic mesh that provided a favorable attachment surface for landing bats. We trained wild-caught bats to land on the platform by positively reinforcing successful landings with food rewards (fruit and juice for frugivorous bats, mealworms for insectivorous bats, and water for all bats), and recorded their landing maneuvers with a synchronized array of three high speed video cameras (Phantom Miro M340, Vision Research, Wayne, NJ, USA; 800 frames per second, 1000 μs exposure; Lenses: Sigma DC 17- 50mm 1:28 EX HSM, SIGMA Corporation, Ronkonkoma, NY, USA) and three LED lights (Veritas Constellation 120, Integrated Design Tools, Pasadena, CA, USA).

Sample sizes for number of species, number of individuals per species, and number of landings per individual were subject to species availability at field sites and the extent to which wild-caught individuals were amenable to training. Previous studies documented extremely low, and in some cases nonexistent, intraspecific variation in landing style (Boerma et al., 2019; Riskin et al., 2009). We therefore accepted samples of one individual per species, but required at least two landings per individual. We trained a subset of 65 individuals (18 species) to land on a ceiling-mounted force plate (ATI nano17, ATI Industrial Automation, Apex, NC, USA fitted with custom acrylic mounting and landing plates). We used a custom MATLAB script to sample impact forces at 1000 Hz, and to synchronize data collection between the force transducer and the high speed cameras using a post-trigger initiated at the end of a landing event.

### Ceiling reaction forces

We filtered the force profiles using a zero-phase 2^nd^ order low-pass Butterworth filter with a cutoff frequency of 100 Hz, which attenuates high-frequency oscillations and electrical noise while preserving the primary peaks associated with landing impact. Although filtering diminishes the absolute magnitude peak forces, accurate comparisons among individuals for all force components are preserved as long as they have been filtered using the same parameters (Boerma et al., 2019; Riskin et al., 2009). Our filtering parameters match those used by previous investigations of bat landing impact forces (Riskin et al., 2009 and Boerma et al., 2019). We normalized landing impact forces to each individual’s bodyweight (BW), calculated from the difference between an unloaded plate just prior to landing and the bat’s hanging weight once landed (mass also verified prior to data collection using a Pesola scale), then extracted peak 3D impact force into the plate for each landing. We averaged peak impact forces for each individual prior to statistical tests.

### Definitions of categorical variables: landing style and roosting ecology

We used high speed videography to categorize bat landings according to the convention established in Riskin et al. 2009, which names landing styles according to the number of limbs that make initial contact at landing impact with the roost. Landing styles include two-point landings (both hind limbs), three-point landings (both hind limbs plus one thumb claw), and four-point landings (both thumb claws plus both hind limbs) (figure1, landing style insets, Supplemental Videos 1 – 5).

We classified the roosting habits of each species according to published observations (table 1), using categories for roosting guilds outlined in Voss et al. 2016 and Garbino & Tavares 2018, with modifications. Our roosting categories included: cavity in standing tree (CST), exposed on standing tree (EST), unmodified foliage (FOL-UF), furled leaf-tubed (FOL-TB), foliage modified into leaf-tents (FOL-LT), termite or ant nests (TAN), rocks and/or caves (R/C), and rock crevices (CREV).

### Phylogenetic analyses: Ancestral state reconstruction, phylogenetic ANOVA, and phylogenetic logistic regression

We used a published time-calibrated molecular phylogeny (Shi and Rabosky, 2015), pruned to our focal taxa, for all phylogenetic analyses (excluding *A. watsoni*, which was not included in the Shi & Rabosky tree), using the Phytools R-package (Revell, 2018, 2011). We then assigned one landing style as a discrete character to each taxon according to its most-often observed landing style (table 1).

We conducted an ancestral state reconstruction using stochastic character mapping (Huelsenbeck et al., 2003), as implemented in the make.simmap function of the R package phytools, to reconstruct the evolutionary history of landing styles among sampled taxa. We used the fitDiscrete function in the R package Geiger (Harmon et al., 2007) to compare the fit of four different models for the transition matrix of the stochastic character mapping procedure: equal rates, symmetric, all rates different, and meristic. The equal rates model yielded the lowest AICc score, thus we selected this model, which gave all state changes equal probability, and computed the posterior probability for each landing style at internal nodes from 1000 simulated stochastic maps.

Next, we used phylogenetic generalized least squares regression (PGLS), implemented in the R function pgls from the package Caper (Orme, 2018) to explore the extent to which landing impact force is predicted by points of contact. Here, we estimated phylogenetic signal using the maximum likelihood value of Pagel’s lambda and treated points of contact and peak 3D landing impact force as continuous variables. We log-transformed impact forces to ensure normality. We then computed a phylogenetic ANOVA (10000 iterations) with post-hoc tests using the phylANOVA function in the R package phytools to test for pairwise differences in log-peak landing impact forces among landing styles. Peak impact force was the response variable and landing style was the factor. We omitted two species from these analyses due to an inability to unambiguously designate them as a two-, three-, or four-point landing: *M. schreibersii* due to its high degree of behavioral variability and *T. tricolor* because it performs a specialized landing maneuver to alight on a vertical substrate (Boerma et al., 2019), rather than beneath a horizonal roost as in the landing experiments for our other sampled taxa.

We used phylogenetic logistic regression with Firth’s correction (Ives and Garland, 2009), as implemented in the R package, *phylolm* (Ho and Ané, 2014), to test the hypothesis that landing styles are associated with the physical properties of roosts. We applied 2000 bootstrap replicates to generate confidence intervals for and test the significance of the model coefficients, ß, which relate to the probability of observing a particular landing style (categorical response variable) given a particular roosting ecology (categorical predictor variables). Positive coefficients indicate a positive association between predictor and response variables, whereas negative coefficients denote a negative relationship. We excluded *T. tricolor* from this analysis because it is the only sampled taxon to perform its landing maneuver and to roost in tubular furled leaves. We compared two models of roosting habits, the latter of which aggregated multiple roost types according to their physical properties, thereby testing our hypothesis that diverse roost types that share physical properties are correlate with landing style. The models were as follows: 1) a null model in which we assigned each taxon’s roosting ecology according to its most-commonly cited roost type (table 1), and 2) an alternate model in which we aggregated roosting ecologies that include stiff, primarily horizontal surfaces (CST, EST, and R/C) into a single category, spatially-constrained, horizontal roosts in vegetation (FOL-LT and TAN) into a second category, compliant horizonal roosts (FOL- UF), and crevices (CREV). We compared the explanatory power of each model using Akaike’s information criterion (AIC).

## Acknowledgements

The authors are grateful for Nancy Simmons, Brock Fenton, Gloriana Chaverri, Rolf Mueller, Holger Goerlitz, Stephan Greif, José Pablo Barrantes, Antonia Hubacheva, Theresa Hügel, and those who attended the annual Belize Batathons for coordinating access to international field sites and assisting in bat capture and experiments. We thank the Max Planck Institute for Ornithology, the Shandong University-Virginia Tech International Laboratory, the Lamanai Outpost Lodge, and the Hacienda Barú Biological Research Station for facilitating our field work. We thank Kenny Breuer and Tom Roberts for providing force transducers, invaluable electronics guidance, and assistance with experiments in the field. We also thank Cosima Schunk, Jorn Cheney, Jeremy Rehm, Andrea Rummel, Lawrence Wang, and Erika Tavares for assistance with experiments and animal husbandry.

## List of Supplemental Files and Source Data Files

- Figure 1–Source Data 1: This .csv file contains the raw peak impact force data, in units of bodyweight, for each recorded landing. Impact forces were recorded at 1000 Hz, normalized to the individual’s body mass, and smoothed using a zero-phase 2^nd^ order low-pass Butterworth filter with a cutoff frequency of 100 Hz, parameters which are identical to those of previous bat landing studies. The total (resultant) force into the ceiling was calculated and the peak extracted for each landing.
- Figure 1–Source Data 2: This .csv file contains the mean peak impact forces for each individual. These values were used to generate Figure 1.
- Figure 1–Source Data 3:
- Figure 2–Source Data 1: This .tree file is the phylogeny from Shi and Rabosky (2015) used for all phylogenetic analyses in this study. The tree was trimmed to include only our focal species prior to any analyses.
- Figure 2–Source Data 2: This .csv file contains the source data for the stochastic character mapping and ancestra state reconstruction for landing style.
- Table 2–Source Data 1: This .csv file contains the source data for the phylogenetic logistic regression results summarized in Table 2.
- PGLS phylANOVA–Source Data 1: This .csv file contains the source data (Taxon, Landing Style, and mean peak impact force) for the phylogenetic generalized least squares regression and phylogenetic ANOVA with pairwise comparisons. This data file omits *Thyroptera tricolor* from both analyses because it is the only species to perform its specialized four-point landing and to roost in furled leaf tubes. It also omits *Miniopterus schreibersii* because its landing style was equivocal. See Supplemental File 3 for a version that includes *Thyroptera tricolor*.
- Supplemental File 1: Posterior probabilities for landing style at each node in the phylogeny shown in Figure 2. Posterior probabilities were estimated using an equal rates model for 1000 simulated stochastic character maps.
- Supplemental File 2: Node legend for posterior probabilities in the phylogeny shown in Figure 2.
- Supplemental File 3: This .csv file summarized the landing style and mean peak impact force for each species for which impacted forces were measured, except *Miniopterus schreibersii*, for which landing style was equivocal. See Table 1, Figure 1, and Source Data files for Figure 1 for impact forces and landing style in *M. schreibersii*.
- Source Code File 1: This .R file contains the code for plotting impact forces shown in Figure 1.
- Source Code File 2: This .R file contains the code for computing the ancestral state reconstruction for landing style using stochastic character mapping.
- Source Code File 3: This .R file contains the code for computing the phylogenetic generalized least squares and ANOVA that test for associations between landing style and landing impact force, and the phylogenetic logistic regressions that test for associations between landing style and roosting ecology.
- Supplemental Video 1: Two-point landing – *Rhinolophus ferrumequinum*
- Supplemental Video 2: Three-point – *Artibeus jamaicensis*
- Supplemental Video 3: Four-point landing – *Sturnira parvidens*
- Supplemental Video 4: Specialized four-point landing – *Thyroptera tricolor*
- Supplemental Video 5: Four-point landing on cave wall – *Myotis myotis*

